# Distinctive tasks of different cyanobacteria and associated bacteria in carbon as well as nitrogen fixation and cycling in a late stage Baltic Sea bloom

**DOI:** 10.1101/775825

**Authors:** Falk Eigemann, Angela Vogts, Maren Voss, Luca Zoccarato, Heide Schulz-Vogt

## Abstract

Cyanobacteria and associated heterotrophic bacteria hold key roles in carbon as well as nitrogen fixation and cycling in the Baltic Sea due to massive cyanobacterial blooms each summer. The species specific activities of different cyanobacterial species as well as the N- and C-exchange of associated heterotrophic bacteria in these processes, however, are widely unknown. Within one time series experiment we tested the cycling in a natural, late stage cyanobacterial bloom by adding ^13^C bi-carbonate and ^15^N_2_, and performed sampling after 10 min, 30 min, 1 h, 6 h and 24 h in order to determine the fixing species as well as the fate of the fixed carbon and nitrogen in the associations. Uptake of ^15^N and ^13^C isotopes by the most abundant cyanobacterial species as well as the most abundant associated heterotrophic bacterial groups was then analysed with a NanoSIMS. Overall, the filamentous, heterocystous species *Dolichospermum* sp., *Nodularia* sp., and *Aphanizomenon* sp. revealed no or erratic uptake of carbon and nitrogen, indicating mostly inactive cells. In contrary, non-heterocystous *Pseudanabaena* sp. dominated the nitrogen and carbon fixation, with uptake rates up to 1.49 ± 0.47 nmol N h^-1^ l^-1^ and 2.55 ± 0.91 nmol C h^-1^ l^-1^. Associated heterotrophic bacteria dominated the subsequent nitrogen cycling with uptake rates up to 1.2 ± 1.93 fmol N h^-1^ cell ^-1^, but were also indicative for fixation of di-nitrogen.

## 1. Introduction

Cyanobacterial mass occurrences are a worldwide phenomenon in limnic, brackish and marine systems. In the Baltic Sea, such blooms occur regularly during summer [1], and due to their high biomasses they significantly add to eutrophication [2,3]. The onset of blooms is promoted by rising water temperatures and low N:P ratios after N-depletion due to the capability of atmospheric nitrogen fixation by several cyanobacterial species [1,3,4]. Total cyanobacterial nitrogen fixation in the Baltic Sea was estimated with 370 kt yr^-1^ [2], and may contribute up to 55% of total nitrogen input [5,6]. Furthermore, filamentous cyanobacteria may contribute for 44% of the community primary production [7]. The major part of nitrogen and carbon fixation is performed in the early summer, followed by a peak in biomass, and ultimately the decay of the bloom in which predominantly recycling processes occur [6,8].

Cyanobacteria as well as eukaryotic phytoplankton live in close associations with heterotrophic bacteria, and interactions between them may range from symbiosis to competition [9,10]. These interactions strongly influence carbon and nutrient cycling and therewith the stability of aquatic food webs [11, 12]. In phytoplankton blooms, heterotrophic bacteria may provide macronutrients via recycling (or fixation) but may be also competitors for inorganic nutrients [11]. Especially at the late stages of cyanobacterial blooms, the associated heterotrophic bacteria may be responsible for a significant share of elemental cycling and fluxes, i.e. for the input of nutrients and organic matter in the ecosystem due to remineralization. Studies on the role of associated bacteria at these late cyanobacterial bloom stages, however, are lacking.

The predominant cyanobacterial genera in Baltic Sea blooms are *Aphanizomenon, Nodularia, Dolichospermum, Pseudanabaena* and *Synechococcus*, whereby the dominant groups and species may differ between years and stage of the bloom [12]. The first three mentioned genera are filamentous and heterocystous, and may form dense surface scums [1]. Baltic Sea *Synechococcus* sp. and *Pseudanabaena* sp. are supposed to be not capable of nitrogen fixation and hence depend on dissolved nitrogen sources [13,14], even though nitrogenase genes occur in *Pseudanabaena* [14,15]. Thus, *Aphanizomenon, Nodularia*, and *Dolichospermum* are thought to dominate the biological nitrogen input into the Baltic Sea [14]. Recently, however, heterotrophic bacteria were shown to be capable of nitrogen fixation at depth in the central Baltic Sea [16] and may even be the principle N_2_ fixing organism in a Baltic Sea estuarine [17]. However, studies that examined carbon and nitrogen fixation in cyanobacterial blooms and associated heterotrophic bacteria mostly focussed on single cyanobacterial species [7,18], or neglected associated bacteria as well as the fate of the fixed carbon and nitrogen in the associations [14]. In the present study, we incubated a natural late stage Baltic Sea cyanobacterial bloom with ^13^C bi-carbonate and ^15^N_2_, and followed the uptake over time by means of NanoSIMS technique. Therewith, we aimed at unravel the specific contribution of different cyanobacterial species and associated heterotrophic bacteria in carbon and nitrogen fixation as well as the fate of the fixed carbon and nitrogen in the associations.

## 2. Material and Methods

### 2.1. Incubation experiments

A natural cyanobacterial bloom was sampled at station TransA (58°43.8‘N, 18°01.9‘E, Fig. 1) on 13.08.2015. Bloom samples were further concentrated by means of a light trap to remove positive phototactic zooplankton until a cyanobacterial chl. a concentration of 9 µg l^-1^ was reached (measured with a PHYTO-PAM, Heinz Walz GmbH). At Askö laboratory (ca. 1 h transfer), five 176 mL opaque Nalgene bottles were filled with the concentrated bloom till overflowing and sealed with septum caps enabling addition and retrieval of liquids with syringes.

**Figure 1:**
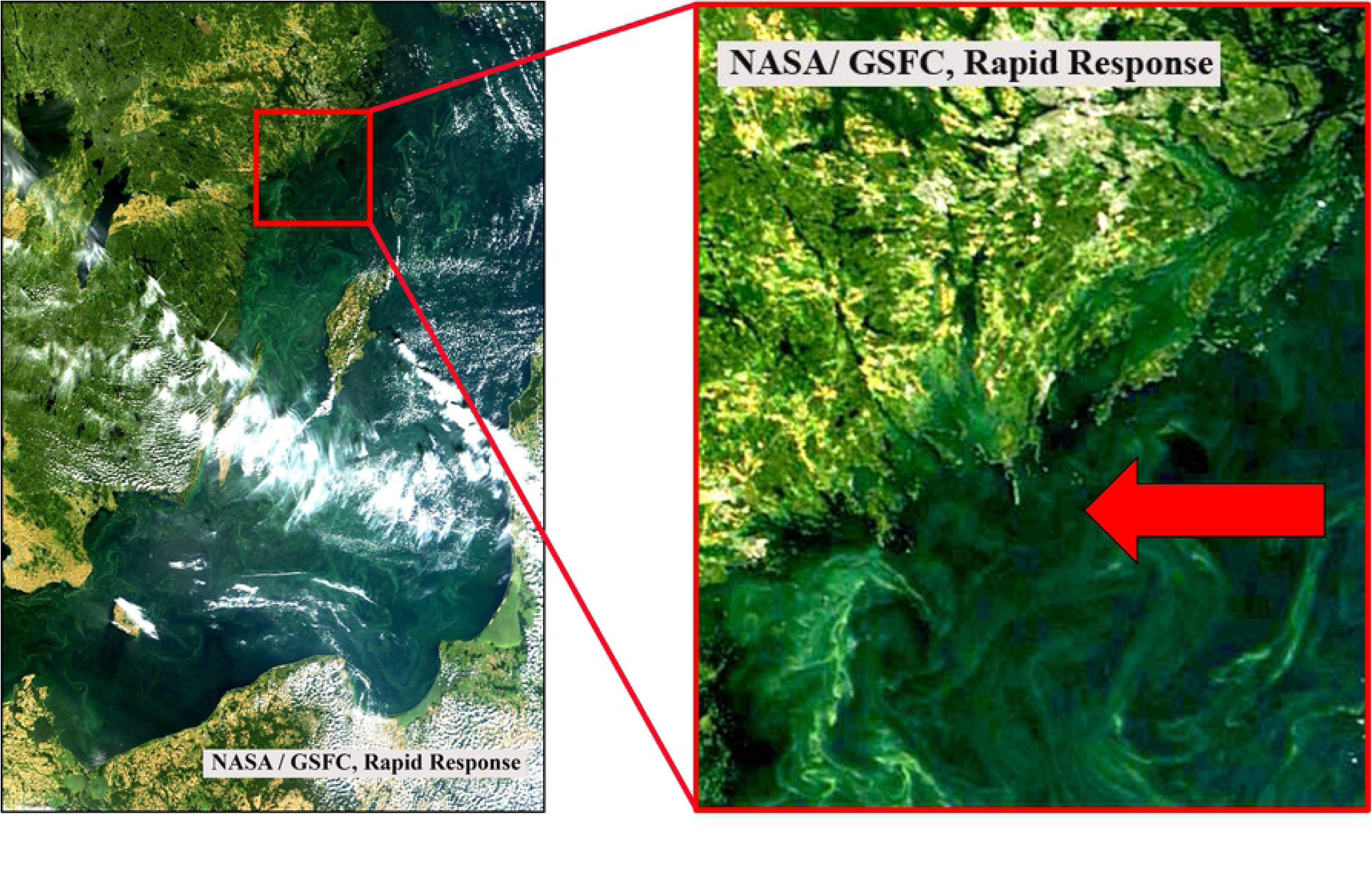
True color satellite image of a cyanobacterial bloom in the Baltic Sea on August 13, 2015 derived from MODIS/Terra (NASA/GSFC, Rapid Response). The arrow in the zoom image on the right side points towards the sampling station TransA.

For ^15^N addition, 1 mL of the sample was removed and subsequently 1 mL 99% pure ^15^N_2_ gas injected with a syringe, resulting in 31.68 atom % ^15^N. For amending ^13^C, 5 ml sample were removed with a syringe and subsequently 5 ml F/2 medium [52] without nitrogen source, adjusted to 8 PSU and spiked with 0.4 g NaH^13^CO_3_ added (final concentration 108.36 atom % ^13^C). Incubation times were 10 min, 30 min, 1 h, 6 h and 24 h. Bottles were incubated in an incubation chamber at 16.5 ± 0.5 °C at approximately 60 µmol photons s^-1^ m^-2^ (delivered from ROHS 36W 840 light bulbs), resembling the natural conditions of sampling under constant light (Supplemental 1).

### 2.2. Sampling

Each sample was fixed with formaldehyde (2% final concentration) for 3 h in the dark at room temperature, and filtered gently onto 3 µm pore width polycarbonate filters for later inspection with CARD-FISH and NanoSIMS. Before start of the incubation, 80 mL of the stock sample were filtered onto 3 µm pore width polycarbonate filter for DNA extraction of the associated bacterial fraction. For phytoplankton counting, a 100 mL subsample was fixed with an acidic Lugol solution [19] and counted according to the Utermöhl technique. To determine biomass percentages, the carbon content (µg l^−1^) of each species was calculated using the official PEG Biovolume Report 2016 (International Council for the Exploration of the Sea) for phytoplankton species and the carbon content per counting unit for the respective size class.

### 2.3. DNA extraction

DNA was extracted as described in [20] with modifications. Briefly, the filters were cut into pieces and mixed with sterilized zirconium beads, 500 μl of phenol/chloroform mix, and 500 μl of SLS extraction buffer. After centrifugation of the mixture, the supernatant was transferred to another tube and the process was repeated. DNA was precipitated overnight at −20°C. The pellet was washed with ethanol, dried, and resolved in autoclaved DEPC-treated water.

### 2.4. PCR and sequencing

For PCRs, 10 ng of DNA was added to autoclaved DEPC-treated water, 10× PCR buffer, BSA, MgCl_2_, dNTPs, forward and reverse primers, and native Taq polymerase. Bacterial DNA was amplified using the primers 341f and 805r [21], under the following conditions: 30 cycles of denaturation for 40 s at 95°C, 40 s of annealing at 53°C, and 1 min of elongation at 72°C. PCR products were cleaned with the Nucleospin kit following the manufacturer’s instructions and shipped to LGC Genomics GmbH (Berlin). Illumina MiSeq V3 sequencing with 300 bp paired-end reads was performed using the 16S primers 341F and 785R. The forward and reverse reads were deposited at the European Nucleotide Archive under the accession number PRJEB23316 (sample B15_3). Taxonomic identification of the associated bacterial community, was performed as described in [22] with the NGS analysis pipeline of the SILVA rRNA gene database project (SILVAngs 1.3).

### 2.5. CARD-FISH analyses

The Illumina runs mostly yielded Alphaproteobacteria and Cytophaga/Bacteroidetes (Fig. 3), and probes Alf968 [23] and CF968 [24] were chosen for analyses of associated heterotrophic bacteria. CARD FISH analyses were computed as described in [25] with modifications: Filter pieces were doused in 0.2 % fluid agarose, dried, and subsequently incubated for 60 min at 37 °C in 10 mg ml^-1^ lysozym solution and thereafter for 15 min at 37 °C with achromopeptidase (180 U ml^-1^). For inactivation, filter pieces were doused subsequently to 1x PBS, autoclaved MilliQ and 99% ethanol and following placed for 10 min in 0.01 M HCl at room temperature. Hybridization with horseradish peroxidase labeled 16S rRNA probes Alf968 and CF968 were carried out at 35 °C with 55% formamide for 3.5 and 4 h, respectively. Signal amplification was achieved with Oregon green 488-X bound to tyramide as described in [26]. After hybridization, filter pieces were stained with 4,6-diamidin-2-phenylindol (DAPI) solution for unspecific counter-staining of all cells.

**Figure 2:**
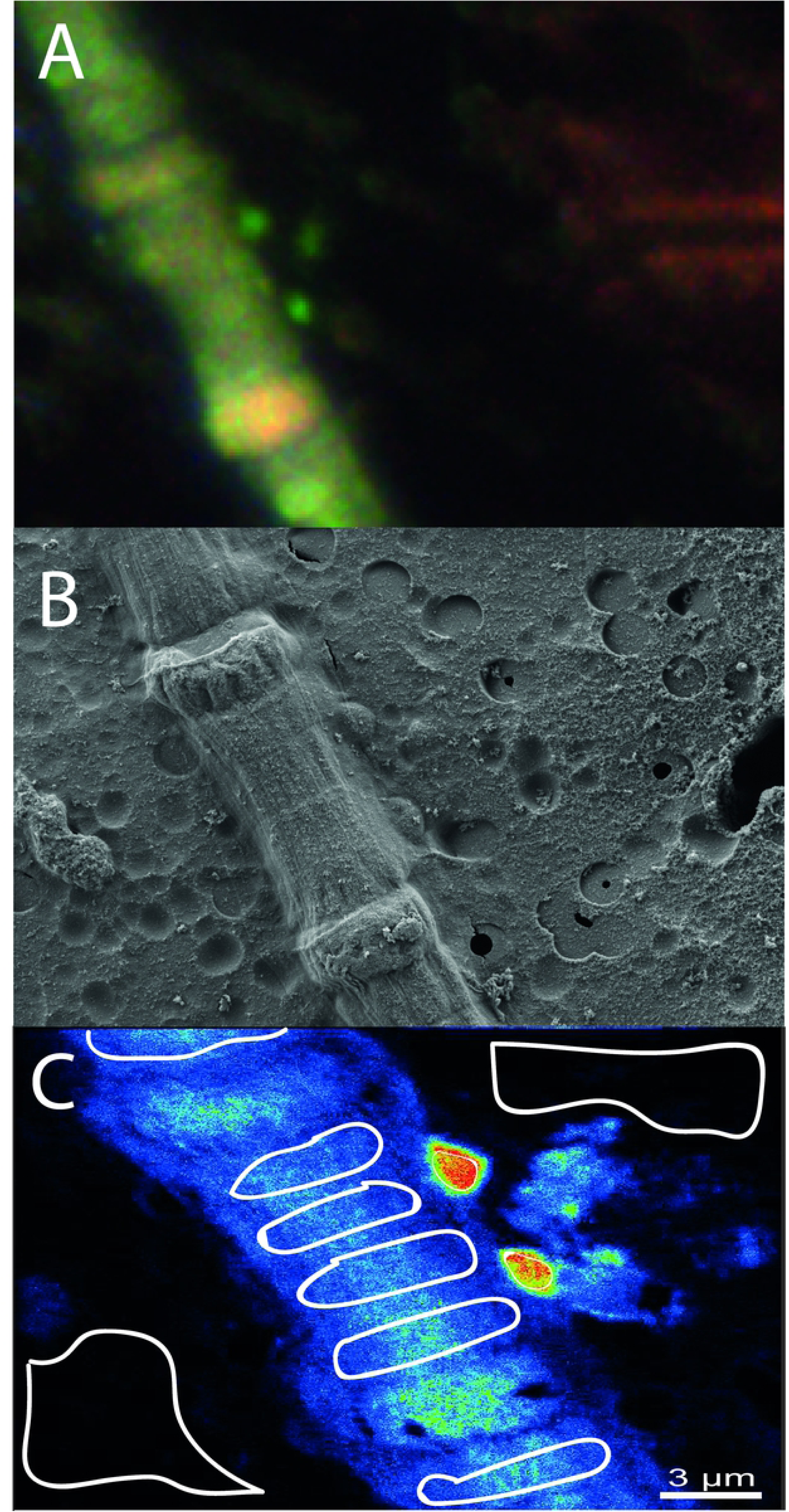
Work flow for analyses of cyanobacteria and their associated heterotrophic bacteria. A: Card-FISH image of a *Nodularia* sp. filament with two associated Alphaproteobacteria taken with a laser microdissectional microscope. The marking arrow can be seen at the right side. B: Scanning electron microscope image of the same spot for confirmation of associated bacteria (middle-right side of the filament) and identification of *Nodularia* sp. The tip of the marking arrow can be seen at the right side of the image. C: accumulated NanoSIMS images of the same spot with blue (low) to red (high) ^15^N signal (as example). The circled areas display the regions of interest, whereof ^13^C/^12^C and ^15^N/^14^N ratios were calculated. Control (filter without cyanobacteria or heterotrophic bacteria) regions can be seen at the top-right and down-left, *Nodularia* sp. regions are displayed in the bluish part, and the associated Alphaproteobacteria by the smaller regions in the reddish part of the image.

**Figure 3.**
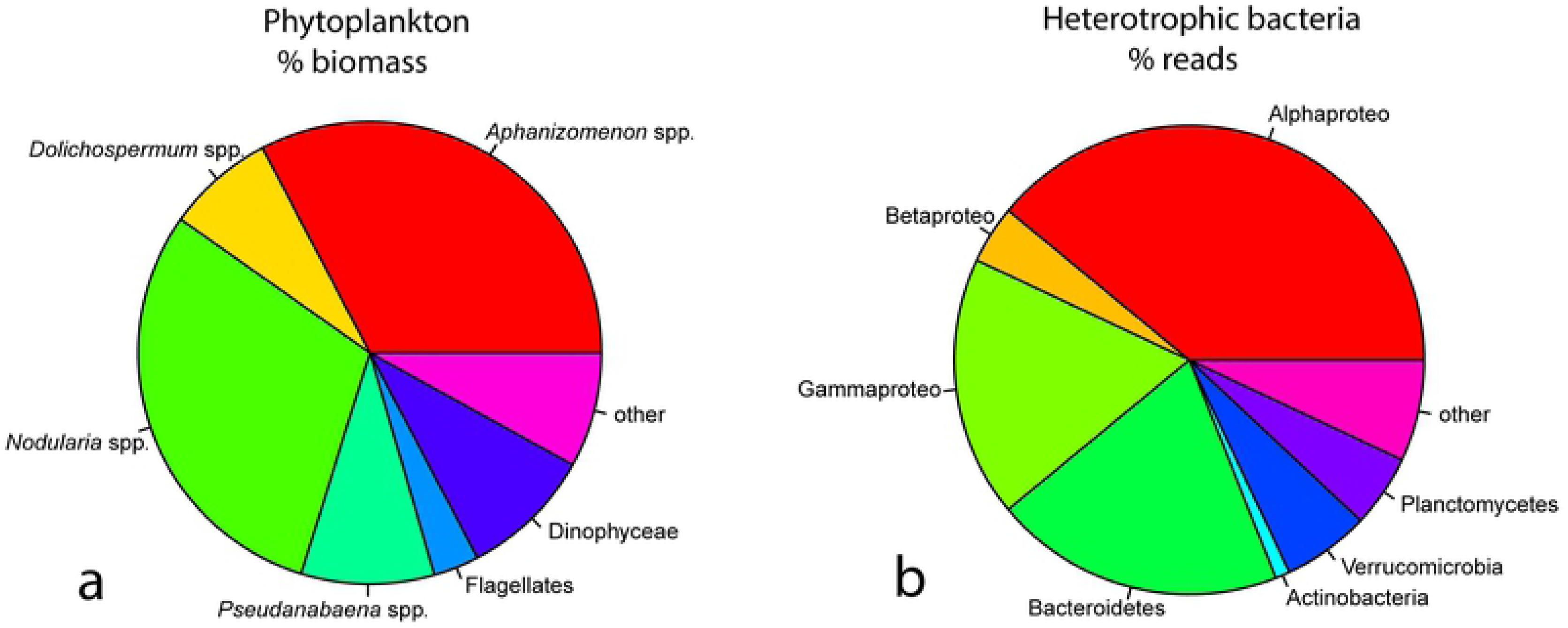
A: Pie chart for the most abundant phytoplankton groups (left side, in % biomass). B: Pie chart for the most abundant bacterial groups (right side, in % of sequencing reads).

### 2.6. Laser-Scanning Microscope, Scanning electron microscope and sputtering

Spots of interest were determined by fluorescence microscopy and subsequently laser marked with a laser microdissectional microscope. For confirmation of associated bacteria and cyanobacterial species, SEM analyses were performed. Therefore, filter pieces were covered with approximately 8 nm gold in a sputter coater (Cressington108 auto-sputter coater). Samples were analyzed with a Scanning electron microscope (Zeiss Merlin VP compact) with the Zeiss Smart SEM Software. Before NanoSIMS analyses, filter pieces were covered with ca. 30 nm additional gold with a sputter coater (see above).

### 2.7. NanoSIMS measurements

SIMS imaging was performed using a NanoSIMS 50L instrument (Cameca, France). A ^133^Cs^+^ primary ion beam was used to erode and ionize atoms of the sample. Among the received secondary ions, images of ^12^C^-, 13^C^-, 12^C^14^N^-^ and ^12^C^15^N^-^ were recorded simultaneously for cells at the laser microdissectional (LMD)-marked spots using electron multipliers as detectors. The mass resolving power was adjusted to suppress interferences at all masses allowing, e.g. the separation of ^12^C^15^N^-^ from interfering ions such as ^13^C^14^N^-^. Prior to the analysis, sample areas of 30×30 µm were sputtered for 2 min with 600 pA to erode the gold and reach the steady state of secondary ion formation. The primary ion beam current during the analysis was 1 pA; the scanning parameters were 512×512 px for areas of 20–30 µm, with a dwell time of 250 µs per pixel.

### 2.8. Analyses of NanoSIMS measurements

All NanoSIMS measurements were analysed with the Matlab based program look@nanosims [27]. Briefly, the 60 measured planes were checked for inconsistencies and all usable planes accumulated, regions of interest (i.e. cells of cyanobacterial filaments, associated bacterial cells and filter regions without organic material for background measurements) defined based on ^12^C^14^N mass pictures, and ^13^C/^12^C as well as ^15^N/^14^N ratios calculated from the ion signals for each region of interest. Measurements of heterocysts in *Aphanizomenon* sp., *Dolichospermum* sp., and *Nodularia* sp. were avoided due their specific cell metabolism. For analyses of each measurement, first the means of background measurements were determined, and this mean factorized for theoretical background values (0.11 for ^13^C/^12^C and 0.00367 for ^15^N/^14^N). These factors were applied to all non-background regions of interest in the same measurement. For each time-point, values for each species (or bacterial group for the associated bacteria) were pooled (i.e. different cells in one measurement as well as different measurements) and means for each species (or bacterial group for the associated bacteria) for each time-point calculated. Work flow for an example spot from Card-FISH to NanoSIMS analyses is illustrated in Fig. 2. The numbers of measured cells per species/group and time point, as well as overall measured areas per time point are given in Supplemental 2.

### 2.9. Uptake rates of ^13^C and ^15^N

Uptake rates for nitrogen and carbon were calculated as described in [28] according to the equation:

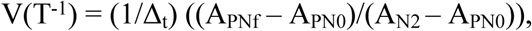

where A_N2_ is the ^15^N or ^13^C enrichment of the N or C available for fixation; A_PN0_ the ^15^N or ^13^C enrichment of particulate N or C at the start of the experiment; A_PNf_, the^15^N or ^13^C enrichment of particulate N or C at the end of the experiment; and

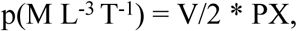

where PX is the concentration of N or C for the respective cyanobacterial species in the incubation bottles, or the cellular N or C content for the associated bacteria. The solubility of N and C was calculated using the Excel Sheet provided by Joe Montoya, based on [29] for CO_2_ and [30] for N_2_. For cyanobacteria gross uptake rates were calculated per volume and time, and for the associated bacteria per cell and time, because no absolute numbers of associated bacteria were existent. The C:N ratios in the cyanobacteria were assumed with 6.3 [14,31]. The size of the associated bacteria was assumed with 2×1 µm (SEM analyses), the carbon content with 0.35 pg C µm^-3^ [32], and the C:N ratio with 5:1 [33]. We are aware that the used “bubble-method” for injection of N_2_ gas assumes an instantaneous equilibrium between the ^15^N_2_ gas bubble and the N_2_ dissolved in water, which in fact may be time-delayed [34], and ultimately leads to an underestimation of fixation rates. Thus, especially at the early measuring points (10 and 30 min), the calculated rates should be considered as proxy values with percentage errors up to 70% [35].

### 2.10. Data analyses

All data were analysed with R studio [36]. To test for differences in stable isotope ratios between species/groups or between different time-points in the same group/species, ANOVAS (analyses of variance) with subsequent Tukey HSD posthoc tests with the package agricolae were performed. Likewise, the impact of the host species on the stable isotope uptake of the associated bacteria was tested with ANOVAs, by comparing associated bacterial cells from different hosts. Possible cell-to-cell transfer of ^13^C and ^15^N between host and associated bacteria were tested by calculating linear models of ^13^C/^12^C and ^15^N/^14^N ratios between the host cells and the associated bacterial cells for each incubation period. To test for correlations between ^13^C and ^15^N uptake, linear models were calculated with the lm function. To test for differences in relations of ^13^C to ^15^N uptake between species/groups, dissimilarity matrices (horn distances) were calculated with a xy (x = ^13^C/^12^C, y = ^15^N/^14^N) system, and subsequently ANOSIM analyses performed with the vegan package. To test for differences in ^13^C/^15^N uptake relations between functional groups, ANCOVAs with and without interactions between the factor and the co-variable were calculated with linear models. Here, ^13^C uptake was set as dependent variable, ^15^N uptake as co-variable, and the functional group as factor. Next, ANOVAs were calculated for both ANCOVAs to test for differences in the slopes of the linear models.

## 3. Results

### 3.1. Community composition of the phytoplankton and associated bacteria

The phytoplankton community was dominated by the cyanobacteria *Aphanizomenon* sp. (33% biomass), *Nodularia* sp. (30% biomass), *Pseudanabaena* sp. (9% biomass) and *Dolichospermum* sp. (8% biomass), which together accounted for 80% of the total biomass (Figure 3a). The most abundant associated bacteria belonged to Alphaproteobacteria (39%), Cytophaga/Bacteroidetes (20%), Gammaproteobacteria (18%), Verrucomicrobia (6%), Planctomycetes (5%), Betaproteobacteria (4%) and Actinobacteria (1%, Fig. 3b).

The general appearance of the bloom (Fig. 4a), and microscopy of cyanobacteria (Fig. 4b-e) both indicated a late stage of the bloom (especially the “curly” appearance of *Nodularia* sp.), with many associated bacteria to the heterocystous species (Fig. 4f).

**Figure 4.**
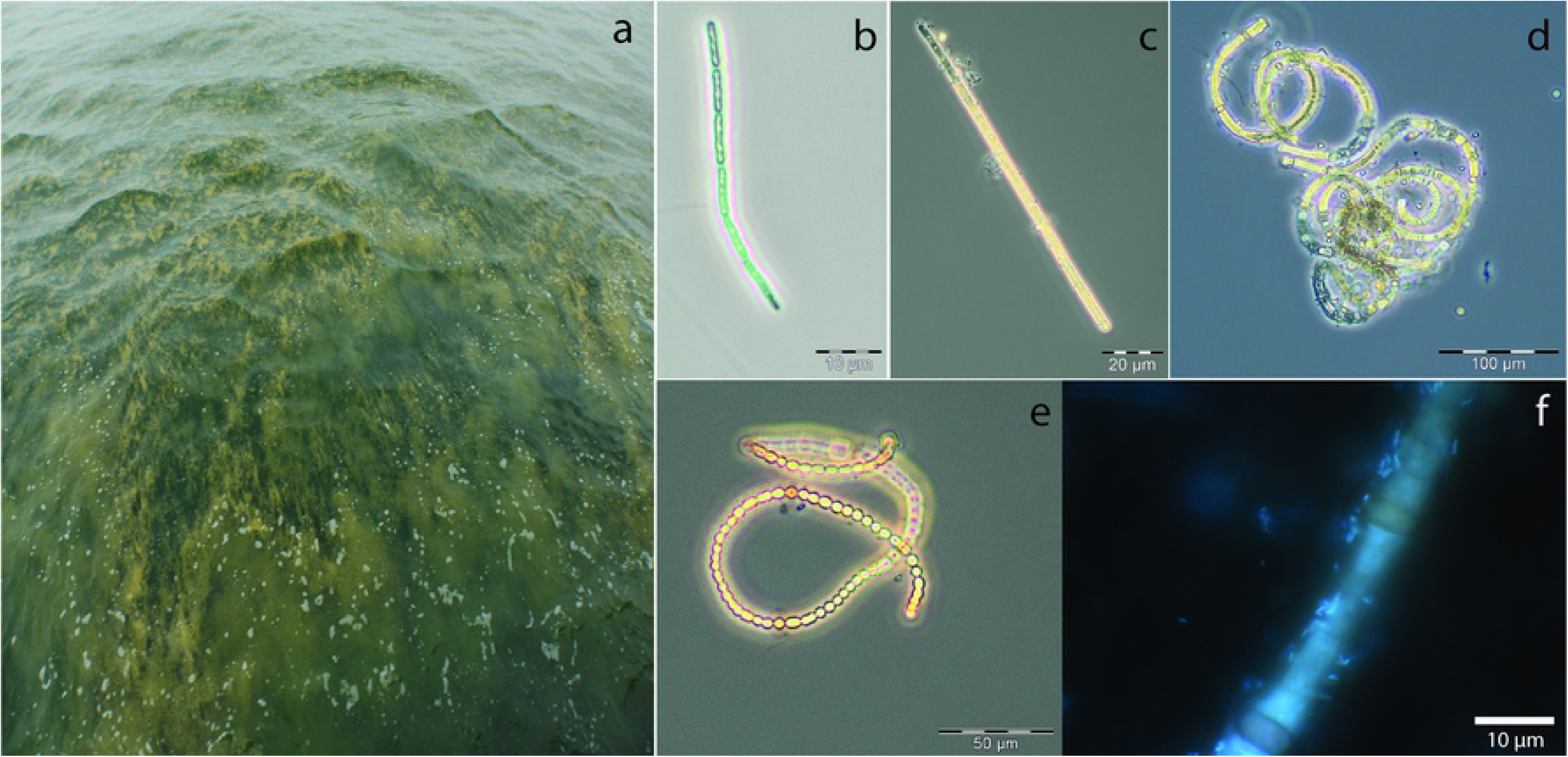
Appearance of the bloom at the day of sampling (a), and microscopic images of *Pseudanabaena* sp. (b), *Aphanizomenon* sp. (c), *Nodularia* sp. (d), *Dolichospermum* sp. (e), and a DAPI stained sample with *Nodularia* sp. and associated bacteria.

### 3.2. Bi-carbonate uptake of cyanobacteria and associated heterotrophic bacteria

Significant differences in the ^13^C incorporation between the bacterial groups were observed at all sampling points (Fig. 5). *Pseudanabaena* sp. showed the highest ^13^C/^12^C ratios at all sampling points with continuously increasing incorporation of ^13^C over time. At the early time points (10, 30 and 60 min), all other species/groups displayed a ^13^C/^12^C ratio close to the natural occurring value of 0.011 (Fig. 5). After 6 and 24 h of incubation, however, Cytophaga/Bacteroidetes revealed the second highest ^13^C/^12^C ratios, corresponding to significant ^13^C enhancements with a more than two- and ten-fold increase of the natural occurring ratio after 6 and 24 h, respectively (Fig. 5). Mentionable, the filamentous cyanobacteria *Aphanizomenon* sp., *Dolichospermum* sp. and *Nodularia* sp. did not display elevated ^13^C/^12^C ratios over the whole 24 h incubation period with two exceptions: *Aphanizomenon* sp. revealed enhanced ratios after 6 h and *Dolichospermum* sp. after 24 h of incubation (data not shown, Fig. 5). To test for a possible impact of the host-species on ^13^C uptake of the associated bacteria, we compared the ^13^C/^12^C ratios obtained from Alphaproteo- and Cytophaga/Bacteroidetes bacteria from different host species. In most cases, however, no significant differences occurred between the hosts (ANOVAs, data not shown). Especially in the 6 and 24 h exposures, where increased ^13^C/^12^C ratios were obtained for both of the associated bacterial groups (Fig. 5), no impact of the host species could be seen (data not shown). Linear models on ^13^C uptake between the host cells and the associated bacterial cells did not suggest cell-cell transfer of ^13^C except for the 60 min incubation (R^2^ = -0.05, 0.12, 0.24, -0.03, -0.05; p = 0.84, 0.24, 0.01, 0.48, 0.75, for 10 min, 30 min, 60 min, 6 h and 24 h incubation, respectively). The calculated uptake rates of the cyanobacteria were highest for *Pseudanabaena* sp. after 60 min with 2.55 ± 0.91 nmol C h^-1^ l^-1^, and from the associated bacteria for Cytophaga/Bacteroidetes bacteria with 0.31 ± 0.34 fmol C h^-1^ cell^-1^ after 24 h of incubation (Table 1).

| *Species/incubation time* | 10 min | 30 min | 60 min | 6 h | 24 h |
| --- | --- | --- | --- | --- | --- |
| *<sup>13</sup>C uptake nmol C h<sup>-1</sup> l<sup>-1</sup> (fmol C h<sup>-1</sup> cell<sup>-1</sup> for associated bacteria)* |  |  |  |  |  |
| *Aphanizomenon* sp. | 0.3 ± 3.52 | 0.00 ± 0.57 | 1.06 ± 2.69 | 0.41 ± 1.35 | 0.00 ± 0.02 |
| *Dolichospermum* sp. | 0.06 ± 0.76 | 0.25 ± 0.71 | 0.05 ± 0.2 | 0.07 ± 0.13 | 0.27 ± 0.18 |
| *Nodularia* sp. | 5.88 ± 20.28 | 0.00 ± 0.95 | 0.4 ± 1.5 | 0.19 ± 0.4 | 0.00 ± 0.03 |

**Table 1:** Carbon and nitrogen uptake rates ± standard deviation given in nmol C or N h^-1^ l^-1^ for cyanobacteria, and fmol C or N h^-1^ cell^-1^ for associated bacteria.

|  |  |  |  |  |  |
| --- | --- | --- | --- | --- | --- |
| *Pseudanabaena* sp. | 2.48 ± 1.5 | 1.98 ± 0.89 | 2.55 ± 0.91 | 1.13 ± 0.72 | 1.87 ± 1.08 |
| *Alphaproteo* | 0.59 ± 0.9 | 0.00 ± 0.04 | 0.00 ± 0.07 | 0.13 ± 0.23 | 0.2 ± 0.32 |
| *Cytophaga/Bacteroidetes* | 0.12 ± 0.09 | 0.24 ± 0.45 | 0.03 ± 0.06 | 0.19 ± 0.27 | 0.31 ± 0.34 |
| *<sup>15</sup>N uptake nmol N h<sup>-1</sup> l<sup>-1</sup> (fmol N h<sup>-1</sup> cell<sup>-1</sup> for associated bacteria)* |  |  |  |  |  |
| *Aphanizomenon* sp. | 1.03 ± 1.6 | 0.00 ± 1.27 | 0.28 ± 0.42 | 0.17 ± 0.4 | 0.00 ± 0.02 |
| *Dolichospermum* sp. | 0.73 ± 0.45 | 0.17 ± 0.3 | 0.05 ± 0.08 | 0.09 ± 0.17 | 0.04 ± 0.03 |
| *Nodularia* sp. | 8.07 ± 18.4 | 0.03 ± 0.87 | 0.18 ± 0.53 | 0.19 ± 0.23 | 0.00 ± 0.02 |
| *Pseudanabaena* sp. | 1.49 ± 0.47 | 0.8 ± 0.56 | 0.84 ± 0.17 | 0.48 ± 0.31 | 0.17 ± 0.05 |
| *Alphaproteo* | 0.00 ± 0.08 | 0.2 ± 0.73 | 0.31 ± 0.76 | 1.15 ± 1.29 | 0.34 ± 0.2 |
| *Cytophaga/Bacteroidetes* | 0.95 ± 1.01 | 0.36 ± 0.53 | 1.2 ± 1.93 | 0.67 ± 0.92 | 0.25 ± 0.17 |

**Figure 5:**
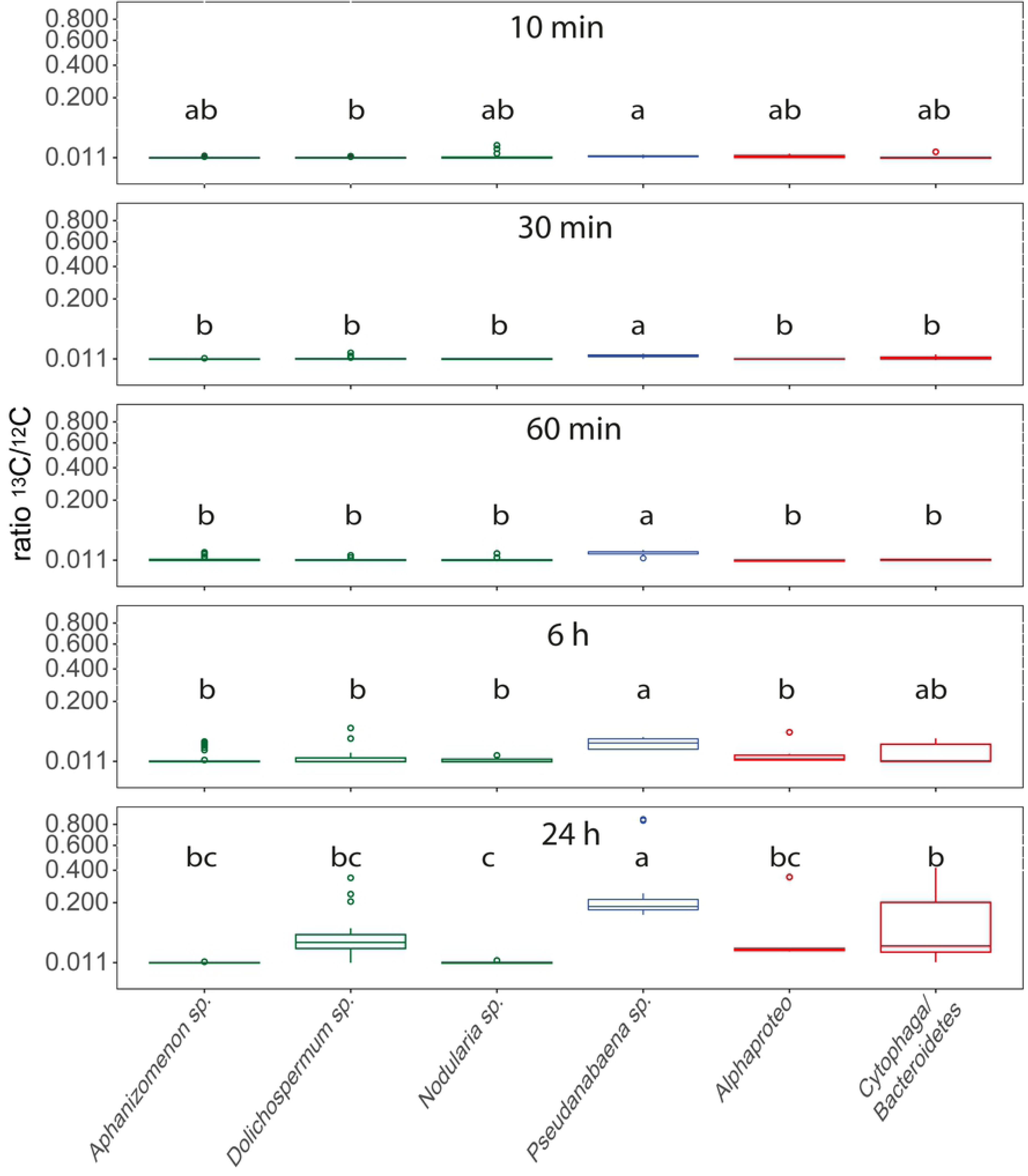
Boxplots of ^13^C/^12^C ratios for *Aphanizomenon* sp., *Dolichospermum* sp., *Nodularia* sp., *Pseudanabaena* sp., Alphaproteobacteria and Cytophaga/Bacteroidetes bacteria over time, with square root transformed y axis. Values originate from pooled data for the respective species from different measurements and cells (Supplemental 2). Lower case letters above the boxplots refer to different groups of Tukey HSD Post-Hoc tests. Heterocystous cyanobacteria are displayed in green, non-heterocystous cyanobacteria in blue, and associated heterotrophic bacteria in red.

### 3.3. ^15^N_2_ uptake of cyanobacteria and associated heterotrophic bacteria

For all time points, significant differences of ^15^N incorporation between the species/groups occurred (Fig. 6). After 10 min of incubation, however, all species still showed values around the natural ^15^N/^14^N ratio of 0.00367 (Fig. 6), whereas after 30 min *Pseudanabaena* sp. (which reveals the highest ^15^N incorporation), and the associated heterotrophic bacteria showed enhanced ^15^N/^14^N ratios (Fig. 6). Between 1 and 6 h of incubation especially the Alphaproteobacteria increased their ^15^N/^14^N ratios, and after 24 h of incubation pronounced differences between the species occured, with associated Alphaproteobacteria showing the highest ^15^N incorporation (mean = 0.0143 ± 0.0059, almost 4-times increased ^15^N/^14^N ratios compared to the natural ratio). In general, after 24 h of incubation the associated bacteria revealed the highest ratios, followed by *Pseudanabaena* sp., whereas the heterocystous cyanobacteria displayed even after 24 h of incubation ^15^N/^14^N ratios close to the natural value (Fig. 6).

**Figure 6:**
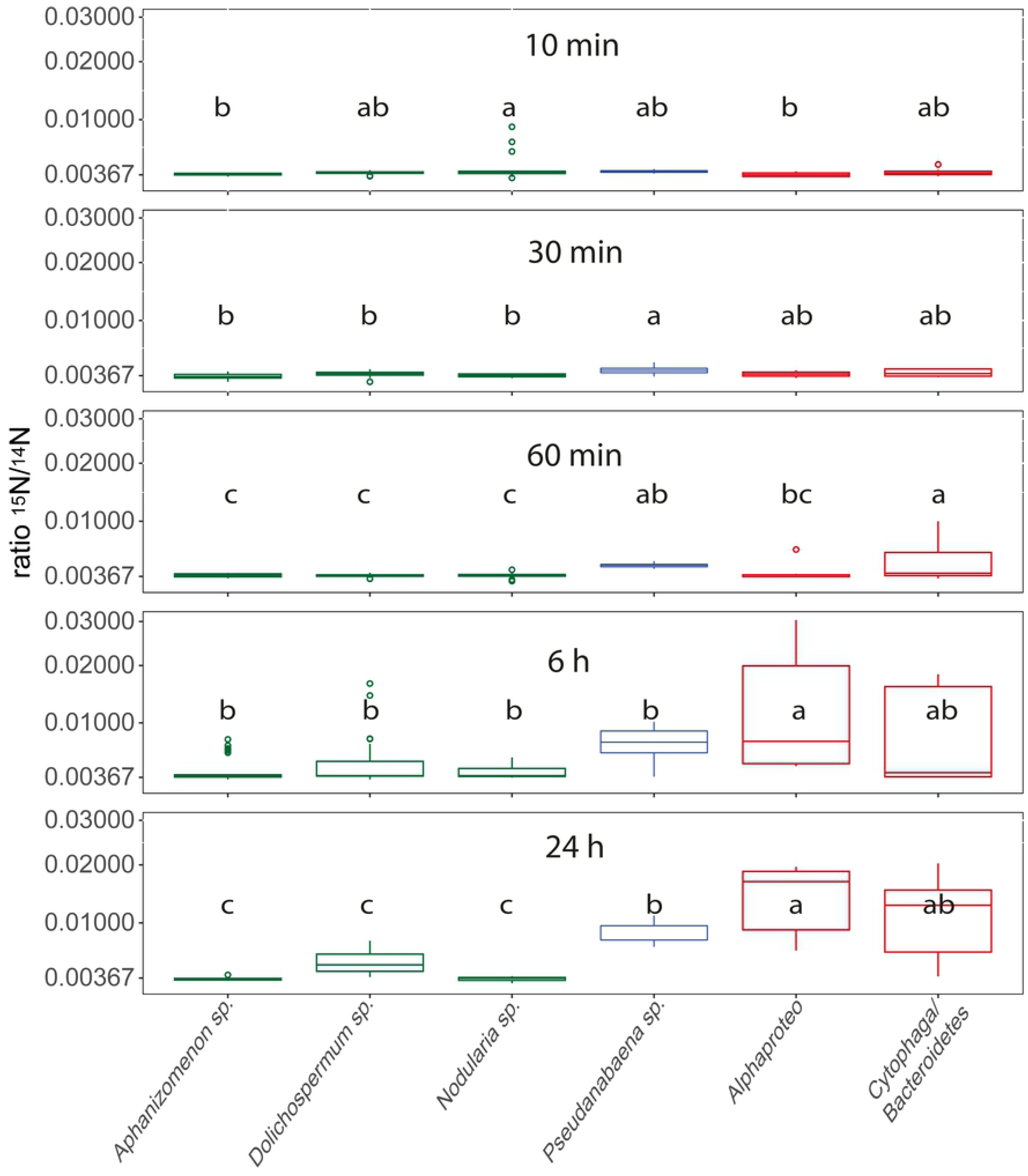
Boxplots of ^15^N/^14^N ratios for *Aphanizomenon* sp., *Dolichospermum* sp., *Nodularia* sp., *Pseudanabaena* sp., Alphaproteobacteria and Cytophaga/Bacteroidetes bacteria over time with square root transformed y-axis. Values originate from pooled data for the respective species from different spots and cells (Supplemental 2). Lower case letters above the boxplots refer to different groups of Tukey HSD Post-Hoc tests. Heterocystous cyanobacteria are displayed in green, non-heterocystous cyanobacteria in blue, and associated heterotrophic bacteria in red.

Comparisons of the ^15^N/^14^N ratios in each species/group between different incubation times revealed significant ^15^N incorporation in most species/groups, but inconsistent ^15^N uptake in the heterocystous species (Fig. 6). In general, the heterocystous cyanobacteria do not display pronounced ^15^N_2_ uptake over time. In contrast, *Pseudanabaena* sp. displays significantly enhanced ^15^N/^14^N ratios from 6 h of incubation onwards (Anova, F = 65.43, p < 0.0001), with steadily increasing values over time and significant differences also between 6 and 24 h of incubation (Fig. 6, Post-Hoc data not shown). Also Alphaproteobacteria and Cytophaga/Bacteroidetes reveal steadily increasing ^15^N/^14^N ratios over the 24 h incubation period (Anova, F = 4.87, p = 0.003, and F = 9.811, p < 0.0001, respectively), with significant differences between almost all incubation times (Fig. 6, Post-Hoc data not shown). The separation of the obtained ^15^N/^14^N values of associated Alphaproteo- and Cytophaga/Bacteroidetes bacteria by the host species did not reveal differences between the host species (data not shown). Linear models between the ^15^N/^14^N ratios of heterocystous cyanobacterial cells that carry associated bacteria and the associated bacteria did not suggest dependencies of ^15^N uptake between the host and the associated bacterium, with the exception of 30 min incubation (R^2^ = -0.02, 0.64, 0.08, -0.03, 0.1; p = 0.48, 0.02, 0.1, 0.39, 0.1 for 10 min, 30 min, 1h, 6 h, and 24 h incubation, respectively). The uptake rates were highest for *Nodularia* sp. with 8.07 ± 18.4 nmol N h^-1^ l^-1^ after 10 min of incubation. However, if excluding the 10 min incubation due to experimental uncertainties, *Pseudanabaena* sp. revealed the highest incorporation rates with 0.84 ± 0.17 nmol N h^-1^ l^-1^ after 1 h of incubation. For the associated bacteria Cytophaga/Bacteroidetes displayed the highest incorporation of ^15^N with 1.2 ± 1.93 fmol N h^-1^ cell^-1^ after 1 h incubation (Table 1).

### 3.4. Species- and group specific relations of ^13^C to ^15^N uptake

Significant differences between the species/bacterial groups occurred for all time points for relations of ^13^C against ^15^N uptake (ANOSIM, each p = 0.001), although different R values were obtained for different exposure times (R = 0.2387, 0.4203, 0.3098, 0.215, 0.585, for 10, 30, 60 min, 6 and 24 h exposure, respectively), indicating most pronounced differences in the relation of ^13^C to ^15^N uptake between the species/groups after 24 h of incubation. In general, *Pseudanabaena* sp. was the most noticeable species in the ^13^C uptake (starting with the 30 min exposure), and the associated Alphaproteo and Cytophaga/Bacteroidetes bacteria in the ^15^N uptake (starting after 60 min of exposure, Fig. 7). The heterocystous cyanobacteria revealed a high patchiness with few cells displaying prominent ^13^C uptake (Fig. 5), but mostly did not show obvious uptake of either ^13^C or ^15^N (Fig. 7, Table 1). Pooling the different species (for bacteria groups) into the functional groups heterocystous cyanobacteria (*Aphanizomenon* sp., *Dolichospermum* sp., and *Nodularia* sp.), non-heterocystous cyanobacteria (*Pseudanabaena* sp.), and associated bacteria (Alphaproteo- and Cytophaga/Bacteroidetes bacteria), and plotting of the ^13^C/^12^C and ^15^N/^14^N ratios against the time, revealed specific tasks of the functional groups (Fig. 7, Table 2). The associated bacteria predominantly display enhanced ^15^N/^14^N ratios, with the highest ratios after 6 h incubation, whereas non-heterocystous cyanobacteria reveal the highest ^13^C/^12^C ratios with a time dependent increase. Controversially, only few heterocystous cyanobacteria show increased ^13^C/^12^C and/or ^15^N/^14^N ratios (Fig. 7).

**Table 2:** Regression analyses of ^13^C over ^15^N uptake for the functional groups heterocystous cyanobacteria (*Aphanizomeno*n sp., *Dolichospermum* sp., *Nodularia* sp.), associated bacteria (Alphaproteo and Cytophaga/Bacteroidetes bacteria), and non-heterocystous cyanobacteria (*Pseudanabaena* sp.) for the different incubation times. Anova results display comparisons of regression slopes of the different functional groups (ANCOVAs with and without interactions between the factor (functional group) and the co-variable (^15^N uptake) were calculated with linear models, with ^13^C uptake set as dependent variable. ANOVAs were then calculated between both ANCOVAs to test for differences in the regression slopes).

| Incubation time | 10 min | 30 min | 60 min | 6 h | 24 h |
| --- | --- | --- | --- | --- | --- |
| *Heterocystous cyanobacteria* | Y=0.001+0.246x,<br>R<sup>2</sup>=0.88,<br>p=0.000 | Y=0.004-0.024x,<br>R<sup>2</sup>=0.00,<br>p=0.308 | Y=0.001+0.03x,<br>R<sup>2</sup>=0.16,<br>p=0.000 | Y=0.002+0.149x,<br>R<sup>2</sup>=0.82,<br>p=0.000 | Y=0.00+0.01x,<br>R<sup>2</sup>=0.56,<br>p=0.000 |
| *Associated bacteria* | Y=0.004+0.018x,<br>R<sup>2</sup>=0.02,<br>p=0.471 | Y=0.003+0.105x,<br>R<sup>2</sup>=0.45,<br>p=0.06 | Y=-0.02+1.94x,<br>R<sup>2</sup>=0.52,<br>p=0.000 | Y=0.004+0.329x,<br>R<sup>2</sup>=0.4,<br>p=0.007 | Y=0.015-0.022x,<br>R<sup>2</sup>=0.24,<br>p=0.02 |
| *Non-heterocystous cyanobacteria* | Y=0.004+0.028x,<br>R<sup>2</sup>=0.04,<br>p=0.169 | Y=0.003+0.105x,<br>R<sup>2</sup>=0.28,<br>p=0.02 | Y=0.004+0.05x,<br>R<sup>2</sup>=0.72,<br>p=0.001 | Y=0.003+0.117x,<br>R<sup>2</sup>=0.82,<br>p=0.000 | Y=0.009-0.002x,<br>R<sup>2</sup>=0.02,<br>p=0.27 |
| *Anova* | F=2.77, p=0.066 | F=5.69, p=0.001 | F=12.9, p=0.000 | F=60.42, p=0.000 | F=9.79, p=0.000 |

**Figure 7:**
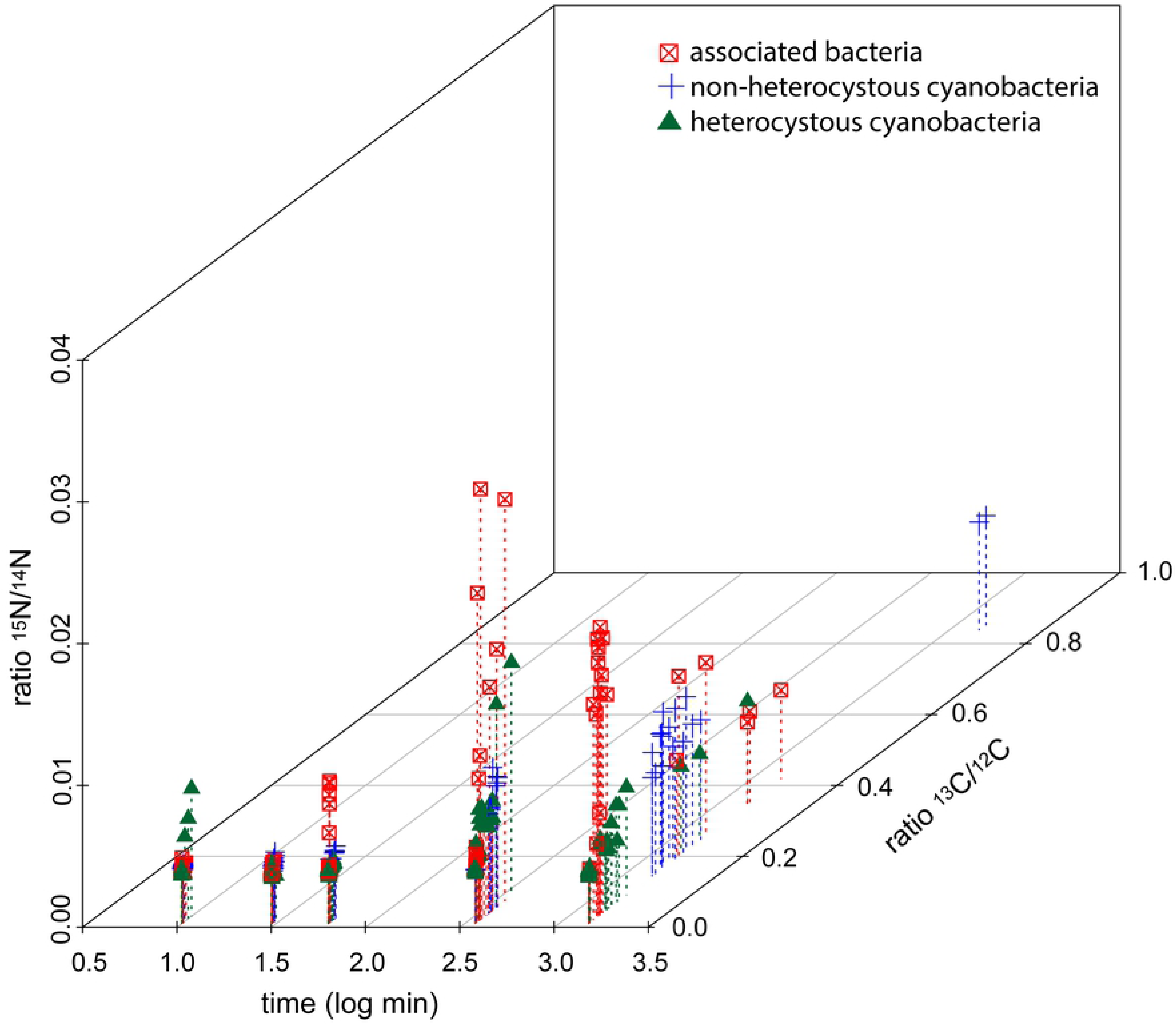
^13^C/^12^C (z axis) and ^15^N/^14^N (y axis) ratios plotted against the exposure time (log transformed x axis) for the different functional groups (heterocystous cyanobacteria, non-heterocystous cyanobacteria, associated bacteria). The color and symbol legend is given directly in the figure.

Group specific behavior was corroborated by significantly different slopes between the functional groups in regression analyses of the ^13^C over ^15^N uptake for the different exposure times, despite the fact that significant correlations between ^13^C and ^15^N uptake occurred for all groups (Table 2). From 60 min of exposure onwards, the slopes of the associated bacteria are by far the steepest, corresponding to a predominant incorporation of ^15^N, whereas non-heterocystous cyanobacteria reveal flat slopes accompanying predominant incorporation of ^13^C (Table 2).

## 4. Discussion

The present study determined the specific contribution of four different cyanobacterial species and the two most abundant associated bacterial groups in carbon as well as nitrogen fixation and cycling in late stage cyanobacterial bloom associations. Altogether, the cyanobacterium *Pseudanabaena* spp. dominated the carbon assimilation as well as nitrogen fixation at the early time-points, and the associated Alphaproteo- and Cytophaga/Bacteroidetes bacteria the nitrogen cycling and possibly N_2_ fixation at the later time-points. The filamentous, heterocystous cyanobacteria *Nodularia* sp., *Dolichospermum* sp., and *Aphanizomenon* sp. on the other hand, either showed no or erratic carbon and nitrogen uptake. Among the associated heterotrophic bacteria Cytophaga/Bacteroidetes were more active in the carbon cycling, whereas Alphaproteobacteria revealed higher activity in nitrogen cycling. However, high intra-species variability was observed in all examined species, which partly impeded significant differences in isotope uptake between species and time points.

### 4.1. Bi-carbonate uptake of cyanobacteria and associated heterotrophic bacteria

Surprisingly, *Pseudanabaena* sp. and not the heterocystous cyanobacteria was the most prominent species in carbon assimilation (Fig. 3), with fixation rates up to 2.55 nmol C h^-1^ l^-1^. Indeed, carbon fixation rates of *Pseudanabaena* sp. were much higher than the rates for the heterocystous species *Aphanizomenon* sp., *Dolichospermum* sp., and *Nodularia* sp. together (Table 1, exception after 10 min of incubation due to 3 extraordinary high measurements in *Nodularia* sp.). However, in combined measurements of June, July and August in the preceding seasons 2012 and 2013, the three heterocystous species together accounted for ca. 5-250 nmol C h^-1^ l^-1^ (Klawonn et al., 2016). Thus, the heterocystous cyanobacteria still hold key roles in carbon fixation in the Baltic Sea [14,37], with much higher fixation rates compared to the estimated ones of *Pseudanabaena* sp. in the present study. In our case, the appearance of the bloom and the curly phenotype of *Nodularia* sp. suggested a late stage of the bloom (Fig. 4), and the low activity of *Aphanizomenon* sp., *Dolichospermum* sp., and *Nodularia* sp. cells might be attributed to inactive cells at the late bloom stage. *Pseudanabaena* sp. was still active and may be adapted to this situation where P-supply by degrading blooms may be granted.

Measurements of heterotrophic bacteria at the later incubation times also revealed enhanced ^13^C/^12^C ratios (Fig. 5), and heterotrophic bacteria may also incorporate bi-carbonate [38]. However, the ^13^C signal in heterotrophic bacteria arises after 6 h incubation which may be related to recycled organic carbon released by *Pseudanabaena* sp. and other cells. The higher proportion of Cytophaga/Bacteroidetes bacteria in the incorporation of ^13^C compared to Alphaproteobacteria (Fig. 5) fits the current knowledge on their ecology: Marine Cytophaga/Bacteroidetes are specialized in the degradation of high molecular weight compounds (Fernández-Gómez et al., 2013; Kirchman, 2002; Alonso et al., 2012), which are especially exuded in high quantities in late stage and senescent blooms (Mühlenbruch et al. 2018; Seymour et al., 2017; Pinhassi et al., 2004). Alphaproteobacteria on the other hand preferentially use low molecular weight compounds such as amino acids [40] and may act complementary to Bacteroidetes/Cytophaga in cyanobacterial bloom associations [39]. Thus, the higher ^13^C incorporation in Cytophaga/Bacteroidetes bacteria may display the recycling of complex organic material whereas the lower signal in the Alphaproteobacteria account for the incorporation of low molecular weight exudates.

### 4.2. ^15^N_2_ uptake of cyanobacteria and associated heterotrophic bacteria

*Pseudanabaena* sp. showed ^15^N_2_ incorporation after 30 min of incubation, and was the only species with significantly increased ^15^N/^14^N ratios at this time. Further, it was the species with the highest ^15^N/^14^N ratios after 60 min of incubation (Fig. 6). Until now, the non-heterocystous *Pseudanabaena* sp. was not shown to be involved in fixation of atmospheric nitrogen in the Baltic Sea [13,14], despite the presence of nitrogenase genes [15]. However, picocyanobacteria and non-heterocystous filamentous species were suspected for nitrogen fixation under specific conditions before [14]. Taking into account that *Pseudanabaena* sp. was the only species with increased ^15^N/^14^N ratios at the early sampling points, our data suggest an active N_2_ fixation by *Pseudanabaena* sp., with fixation rates between 0.17 and 1.49 nmol N h^-1^ l^-1^ (Table 1). Thus, at this late stage of the bloom, *Pseudanabaena* sp. might have been responsible for the input of reactive nitrogen in the multi-species associations and ultimately into the nitrogen cycle of the Baltic Sea. Indeed, if converted to per cell rates, nitrogen fixation of *Pseudanabaena* sp. appears low with up to 0.07 fmol N cell ^-1^ h^-1^ if compared to the heterocystous species *Nodularia spumigena* (11 fmol N cell^-1^ h^-1^, Ploug et al., 2011) and *Aphanizomenon* sp. (1-11 fmol N cell^-1^ h^-1^, Ploug et al., 2010). However, this difference might be attributed to the much smaller cell size of *Pseudanabaena* sp., and compensated by higher cell numbers. In a comparably study of a Baltic Sea cyanobacterial bloom, cumulative fixation rates for combined measurements of June, July and August of the heterocystous species *Dolichospermum* sp., *Nodularia* sp., and *Aphanizomenon* sp. were determined with ca. 0.5-80 nmol N l^-1^ h^-1^ [14], i.e. approximately one dimension above that of *Pseudanabaena* sp. alone in the present study. Likewise to the carbon fixation, the inexistent nitrogen fixation of the heterocystous species in our study may be attributed to different stages of the blooms, with most cells of heterocystous species being inactive at the late stage of the bloom (Figs. 5 and 6). Congenial to these results, early/mid-summer nitrogen fixation rates in the Baltic Sea were up to 30 times higher compared to late summer [8]. Thus, heterocystous cyanobacteria may still be the prime nitrogen fixers in the Baltic Sea [5,6], but the possible participation of *Pseudanabaena* spp. should not be neglected. If this temporal divided nitrogen fixation between different cyanobacterial species represents a general feature for the Baltic Sea needs to be investigated in consecutive studies.

The overall highest ^15^N/^14^N ratios by the associated bacteria after 6 and 24 h of exposure are surprising, taking the high abundance of diazotrophic cyanobacteria and the low ^15^N incorporation of the hosts into account. Indeed, one would expect the converse role allocation, where associated heterotrophic bacteria reveal lower ^15^N/^14^N ratios than their diazotrophic hosts (Adam et al., 2016; Ploug et al., 2011). However, our high ^15^N/^14^N ratios were obtained after 6 and 24 h of incubation, and thus, similar to the ^13^C incorporation, the associated bacteria may have used recycled nitrogen that was originally fixed by cyanobacteria. Supporting this assumption, heterotrophic microorganisms in cyanobacterial associations dominated by *Aphanizomenon* sp. relied on recycled nitrogen [43], and *Aphanizomenon* sp. was shown to release up to 35% of the fixed nitrogen as NH ^+^ [7]. However, direct cell to cell transmission between hosts and associated bacteria was not indicated (see 3.2 and the linear models), and release and transfer of newly fixed N_2_ was not indicated at a similar experiment during 12 h of incubation [14].

The role of heterotrophic bacteria in nitrogen fixation budgets for aquatic ecosystem were recently brought into focus (Bentzon-Tilia et al., 2015; Farnelid et al., 2013), and might have been underestimated in preceding studies [44–46]. As examples, heterotrophic organisms dominated the nitrogen fixation in the South Pacific Gyre [45], and were also the principle nitrogen fixing organisms in a Baltic Sea estuary [17]. Indeed, there are hints that the associated bacteria in our study also performed nitrogen fixation themselves and not only used nitrogen released from other cells: First, if the associated bacteria would only recycle nitrogen that was fixed by other organisms, one would expect a dilution in the ^15^N/^14^N ratios from the primary fixer to the secondary user [8,14,43], which is not the case (Figs. 6 and 7). Second, already after 30 min heterotrophic bacterial cells possessed the overall highest ^15^N/^14^N ratios (Fig. 7), and this fast incorporation indicates active nitrogen fixation. Third, many Alphaproteobacteria (Delmont et al., 2018; Bentzon-Tilia et al., 2015; Farnelid et al., 2013) and Cytophaga/Bacteroidetes bacteria [47] possess nitrogenase genes, and are capable of nitrogen fixation. To validate heterotrophic nitrogen fixation we performed a gene functional analysis with the 16S data of the associated bacteria using paprica - PAthway PRediction by phylogenetIC plAcement [48]. In this analysis, however, only 1.2% of the associated bacteria yielded the full pathway (via ferredoxin) for nitrogen fixation (Supplemental 3) which does not support our assumption. Nevertheless, ecosystem key functions may be performed by low abundant bacteria [49,50], and the per cell fixation rates of the associated bacteria were more than one dimension higher compared to *Pseudanabaena* sp. (1.2 vs 0.07 fmol N h^-1^ cell^-1^), and in the same dimension as uptake rates for the much bigger heterocystous cyanobacteria (0.1 – 32.7 fmol N cell^-1^ h^-1^) in the Baltic Sea [14]. Thus, given the high abundances of associated bacteria, heterotrophic nitrogen fixation might contribute significantly to bulk fixation at this late stage bloom. At this stage of the bloom, senescent phytoplankton exhibit high exudation and leaking rates (e.g. Mühlenbruch et al., 2018), and create an environment with high levels of labile DOC that fuels heterotrophic nitrogen fixation [44,52,53]. This is corroborated with the linear models, where bacteria associated to inactive, senescent hosts showed the highest ^15^N uptake (data not shown). However, until now prerequisites and regulation of heterotrophic nitrogen fixation as well as principle contradictions as fixation in oxygenated waters and at high nitrate and ammonium concentrations are poorly understood [44], and should move into the focus of upcoming studies.

### 4.3. Relation of ^13^C to ^15^N uptake

Significant correlations between ^13^C and ^15^N uptake occurred in most species and at most time points (Table 2), which is in accordance with similar studies from cyanobacterial blooms in the Baltic Sea (e.g. Klawonn et al., 2016). Nevertheless, relations between carbon and nitrogen uptake indicated specific tasks of functional groups (Fig 7, Table 2). *Pseudanabaena* sp. (non-hetercystous cyanobacterium) clearly dominated the ^13^C uptake (Fig. 5) throughout the whole incubation period, but was also the first species with increased ^15^N signals (Fig. 6). For the ^15^N/^14^N ratios, however, *Pseudanabaena* sp. was outpaced by the associated bacteria from 6 h incubation onwards (Fig. 6), and revealed much lower per cell fixation rates (see above). Thus, the associated bacteria may have dominated the nitrogen cycling and possibly fixation at the later sampling points. This specification of functional groups was corroborated by significant different slopes in linear models calculated for correlations between ^13^C and ^15^N uptake (Table 2). The formation of distinct functional groups by different species in late stage bloom associations may ultimately result in the allocation of desired metabolic pathways to every member in the association, including members unable to perform these tasks [54,55]. The concerted action of diverse ecological functions by different functional groups was also proposed for a chlorophyte and its prokaryotic epiflora [56], and might be a general feature of multi-species associations.

## Acknowledgements

This work was supported by the Leibniz Association and a grant of the Human Frontiers Science Program (RGP0020/2016). We thank Annett Grütmüller for NanoSIMS measurements, the field station Askö laboratory for provision of lab space and permission of experiments, Susanne Busch and Regina Hansen for phytoplankton counting, and Joe Montoya for the provision of the Excel Sheet for calculations of uptake rates. The SIMS instrument was funded by the German Federal Ministry of Education and Research (BMBF), grant identifier 03F0626A.

